# Behavioral detectability of optogenetic stimulation of inferior temporal cortex varies with the size of concurrently viewed objects

**DOI:** 10.1101/2022.11.04.514791

**Authors:** Rosa Lafer-Sousa, Karen Wang, Reza Azadi, Emily Lopez, Simon Bohn, Arash Afraz

**Affiliations:** Laboratory of Neuropsychology, National Institute of Mental Health, NIH, Bethesda, MD 20892, USA; Department of Psychology, University of Pennsylvania, Philadelphia, PA 19104

## Abstract

We have previously demonstrated that macaque monkeys can behaviorally detect a subtle optogenetic impulse delivered to their inferior temporal (IT) cortex. We have also shown that the ability to detect the cortical stimulation impulse varies depending on some characteristics of the visual images viewed at the time of brain stimulation, revealing the visual nature of the perceptual events induced by stimulation of the IT cortex. Here we systematically studied the effect of the size of viewed objects on behavioral detectability of optogenetic stimulation of the central IT cortex. Surprisingly, we found that behavioral detection of the same optogenetic impulse highly varies with the size of the viewed object images. Reduction of the object size in four steps from 8 to 1 degree of visual angle significantly decreased detection performance. These results show that identical stimulation impulses delivered to the same neural population induce variable perceptual events depending on the mere size of the objects viewed at the time of brain stimulation.

## Introduction

Artificial perturbation of neural activity in the visual system is known to alter visual perception, presenting a valuable opportunity to explore the causal relationship between neural activity and visual perception. Understanding the nature of the perceptual events evoked by neural perturbations and how they are constrained by the neural state of the visual system is essential for advancing our understanding of the neural mechanisms that underlie vision as a behavior and is essential for identifying the neural underpinnings of visual delusions in psychiatric disease and developing effective visual prosthetics for patients with severe visual impairment. While anecdotal human studies provide a tantalizing glimpse into the potential perceptual effects of cortical stimulation (Brindley and Lewin 1968; Puce, Allison, and McCarthy 1999; Lee et al. 2000; Murphey et al. 2009; Parvizi et al. 2013; Rangarajan et al. 2014; Schalk et al. 2017), high throughput and systematic survey of the case necessitates the appeal to nonhuman primate research.

We investigate the effect of optogenetic stimulation on visual perception in macaque monkeys using Opto-Array (Blackrock Microsystems), a novel chronically implantable array of LEDs, allowing spatiotemporally precise stimulation of the same cortical sites across many sessions (Rajalingham et al. 2021), and a Cortical Perturbation Detection (CPD) task which is highly sensitive (Murphey and Maunsell 2007; May et al. 2014; Dai, Brooks, and Sheinberg 2014; Azadi, Bohn, Lopez, et al. 2022) and unrestricted by prior assumptions about the tuning properties of the targeted neurons or the potential effects of perturbation. Monkeys are trained to detect optogenetic stimulations delivered to regions of their inferior temporal cortex (IT) transduced with the depolarizing opsin C1V1 while viewing images unrelated to the detection task. While behavioral effects of optogenetic manipulation in nonhuman primates have been documented before (Jazayeri, Lindbloom-Brown, and Horwitz 2012; Gerits et al. 2012; Cavanaugh et al. 2012; Ohayon et al. 2013; May et al. 2014; Dai, Brooks, and Sheinberg 2014; Inoue, Takada, and Matsumoto 2015; Afraz, Boyden, and DiCarlo 2015; Acker et al. 2016; Stauffer et al. 2016; El-Shamayleh et al. 2017; Fetsch et al. 2018; Rajalingham et al. 2021), optogenetic studies in monkeys typically produce small behavioral effects (Tremblay et al. 2020; Bliss-Moreau, Costa, and Baxter 2022). The few exceptions utilize tasks that include a behavioral detection component (May et al. 2014; Dai, Brooks, and Sheinberg 2014; Azadi, Bohn, Lopez, et al. 2022). The high sensitivity of the CPD task for optogenetic stimulation has also been supported by rodent studies (Huber et al. 2008); (Histed and Maunsell 2014), and for microstimulation in primates (Murphey and Maunsell 2007; Murphey et al. 2009). While the CPD task has been used in previous optogenetic and microstimulation studies, the novelty of our approach is that we provide a concurrent sensory input (images on a screen) and the content of that input is independent of whether cortical stimulation will occur on a given trial. This enables us to systematically assess the effect of different visual input characteristics on the perceptual events evoked by cortical stimulation.

With this approach we have found that detectability of the perceptual events evoked by optogenetic stimulation of macaque IT cortex highly depends on the content and visibility of the concurrent visual input (Azadi, Bohn, Lopez, et al. 2022). Specifically, our prior study showed that when the monkeys were looking at particular images (e.g. a laundry machine) their ability to detect cortical stimulation was significantly higher than when looking at other images (e.g. a gas mask). Why should looking at one image enhance stimulation detectability and another hinder? Perhaps stimulation adds a consistent visual element (e.g. phosphene, ‘facephene’) to the ongoing contents of perception, independent of the visual input, but the ability to detect it varies across images due to figure-ground effects like crowding or contrast. Alternatively, stimulation may distort the contents of the visual-input driven percept and some distortions are more noticeable, or some images put the visual system in a state that is more or less resistant to the effects of stimulation, leading to perceptual events of different magnitude. That is, does stimulation evoke a consistent perceptual event, independent of the visual input, *detection* of which interacts with the image on the screen? Or, is the *event* induced by stimulation image-dependent in nature?

To tease apart these two interpretations, our prior report tested how diminishing the visibility of the visual input by reducing the contrast, saturation, and spatial frequency of the concurrently fixated images affected detection of the cortical event. If cortical stimulation induces a consistent perceptual event, it should be similarly if not more easily detected when the onscreen images are less visible. However, the study found that stimulation detectability decreased as the visibility of the onscreen images was diminished. This counterintuitive result suggests that the *perceptual nature of the event* induced by stimulation depends on the concurrent visual input. The finding carries significant implications for the design and interpretation of perturbation studies, and for the development of visual prosthetics (*see Discussion*). Given the counterintuitive nature and significance of this finding, we sought to replicate the result using a different kind of visibility manipulation.

In this short report we present a variation on the visibility experiment, diminishing the visibility of the concurrently fixated object images by shrinking them in size, rather than fading and blurring them to uniform gray. While the degree of visual angle subtended by the images will be progressively reduced, their contrast, saturation, and features will be preserved until there is no longer an image on the screen. If the magnitude of the perceptual event evoked by stimulation highly depends on the existence and prominence of the visual input, reducing image size should attenuate stimulation detectability.

## Methods

### Surgical Procedures

In a sterile surgical setting under general anesthesia, two male macaque monkeys (“Ph” and “Sp”) were surgically implanted with Opto-Array (Blackrock Microsystems), a novel chronically implantable 5 x 5 mm array of micro LEDs, over regions of their central IT cortex (on the lateral convexity) transduced with the depolarizing opsin C1V1 (Figure 1A,B). The tissue was transduced by injecting a total of 160 μL of AAV5-CaMKIIa-C1V1(t/t)-EYFP (nominal titer: 8×10^12^ particles/ml) across 16 evenly spaced sites within central IT (10 μl/site), which resulted in a region of ~6mm x 6mm viral expression (left hemisphere in Sp, right hemisphere in Ph; Figure 1B). See detailed surgical methods published elsewhere: (Rajalingham et al. 2021; Azadi, Bohn, Lopez, et al. 2022; Azadi, Bohn, Eldridge, et al. 2022). Arrays were also implanted over the corresponding region of IT cortex in the opposite hemisphere not transduced, to serve as a control site in an earlier study (Azadi, Bohn, Lopez, et al. 2022). All procedures were conducted in accordance with the guidelines of the National Institute of Mental Health Animal Use and Care Committee.

**Figure 1.**
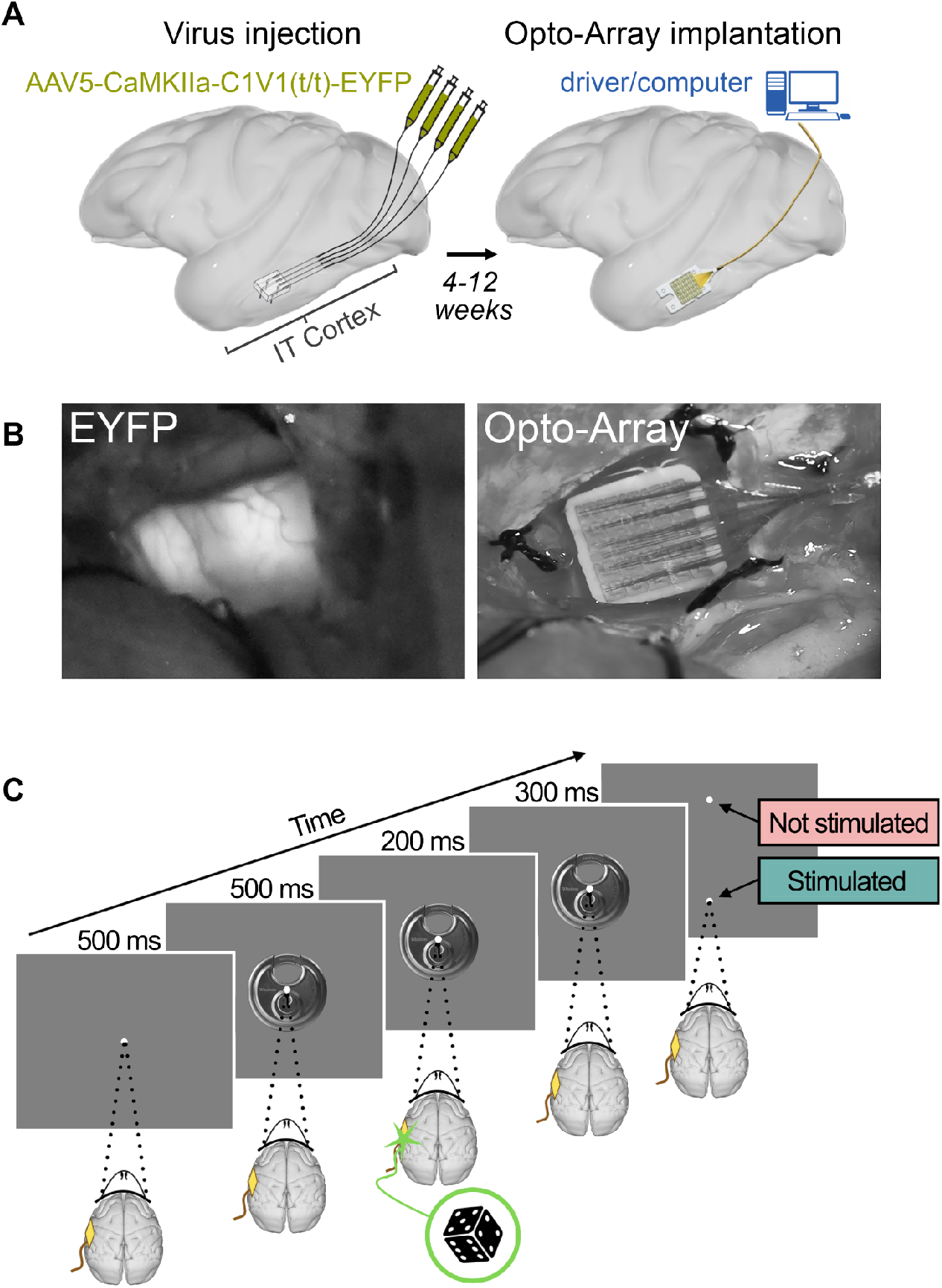
Surgical procedures, task paradigm. **a)** Schematic illustration of the two-part surgical procedure for chronic optogenetic stimulation of IT cortex. Left: Injection of virus AAV5 which expresses the depolarizing opsin C1V1 over a region of central IT cortex in one hemisphere. Right: Implantation of Opto-Array: in a second surgery an Opto-Array was implanted over the transduced region. A control array was also implanted in the corresponding region of central IT in the opposite hemisphere not transduced with virus (control site, not shown). **b)** Confirmation of viral expression (monkey Sp). Left: the signature glow of the fluorescent EYFP protein co-expressed with opsin C1V1. Right: the implanted Opto-Array over the expression zone. **c)** Stimulation-detection task paradigm. Following a 500ms fixation, an image was displayed on the screen for 1s. Halfway through image presentation, a 200ms optical impulse was delivered to the cortex in half of the trials, randomly selected. The animal was rewarded for correctly identifying whether the trial did or did not contain optogenetic stimulation by making a saccade to one of the two points presented at the end of the trial.

### Apparatus

The experiment was carried out with the monkey head fixed, positioned 52 cm from a calibrated 32 in, 120 Hz, 1920×1080 IPS LCD Display++ monitor (Cambridge Research Systems). Fluorescent room lights were turned on. Temperature on the LED die was monitored by a thermistor inside the Opto-Array at the beginning of each trial and trial delivery was paused if the temperature on the LED die rose more than 3° C above the baseline temperature, and restarted once it was less than 1° C above the baseline. A 3° C change at the LED die translates to approximately 0.5° C temperature change on the cortical surface (Rajalingham et al. 2021). A custom MWorks script (The MWorks Project), running on a Mac Pro 2018, was used to control the experiment. A Blackrock LED Driver (Blackrock Microsystems) running a custom firmware version for compatibility with MWorks was used to control the Opto-Array. Gaze was tracked with an Eyelink 1000 Plus (SR Research). Animals received liquid rewards for successfully completing trials and were water-restricted in their cages.

### Stimulation-Detection Task

The monkeys were trained to detect short pulses of illumination delivered to a ~1mm^3^ region of the transduced cortical area (Rajalingham et al. 2021) while viewing images of objects unrelated to the detection task (Figure 1C). The animals were required to maintain fixation within a central 2 degree window throughout each trial (fixation point was 0.15 degrees in diameter). In each trial, following fixation (500 ms), an image was displayed on the screen for 1 s. In half of the trials, randomly selected, optogenetic stimulation was delivered for 200 ms half way through image presentation, and the animal was rewarded for correctly identifying whether the trial did or did not contain cortical stimulation by making a saccade to one of two choice targets presented above and below the screen center Sp was trained to report stimulation trials with a saccade to the upper target, Ph was trained to report stimulation trials with a saccade to the lower target. For each incorrect response, a 3.5 s timeout was enforced. To reinforce the training process, a tone was played when a trial was initiated (following 500ms central fixation), and a second tone was played when the trial completed indicating whether the animal’s response was correct (high pitched tone for correct response, low tone for incorrect response). We previously showed that while the monkeys could learn to accurately report illumination of the arrays implanted over the transduced region, they could not do the same for illuminations at the control site (Azadi, Bohn, Lopez, et al. 2022). For the present report, illumination power was 1.2 mW and 0.81 mW, respectively for Ph and Sp. Due to the sensitivity of the task, the animals can easily get into a saturated level of performance. In order to avoid the ceiling effect, prior to the experiment we determined the illumination levels for each monkey to get them below the ceiling. One experimental session was performed for each monkey with 1493 and 1004 trials collected with overall performance of 80.5% and 87.5% correct trials, respectively for Ph and Sp.

### Stimuli

Each animal performed the stimulation-detection task with an imageset containing 5 object images and 4 size conditions (subtending 1, 2, 4, and 8 degrees of visual angle), plus one “no image” (uniform gray) condition (see Figure 2 for each animal’s image set). The “no image” condition occurred as often as any one image size condition, creating 21 total conditions. In a previous study we had measured the animals’ performance on a battery of 40 images (subtending 8 degrees of visual angle), at two cortical sites (LEDs within the same array) each. For the present study we selected each animals’ top five performing images for a given cortical site. For one animal the exact LED used in the previous study had burned out, so we used a neighboring LED. We have previously shown that the performance image-dependencies are highly correlated for neighboring cortical positions (Azadi, Bohn, Lopez, et al. 2022). That is, we chose each animals’ highest performing images for the present study. Note, the face image in Ph’s imageset is computer generated.

**Figure 2.**
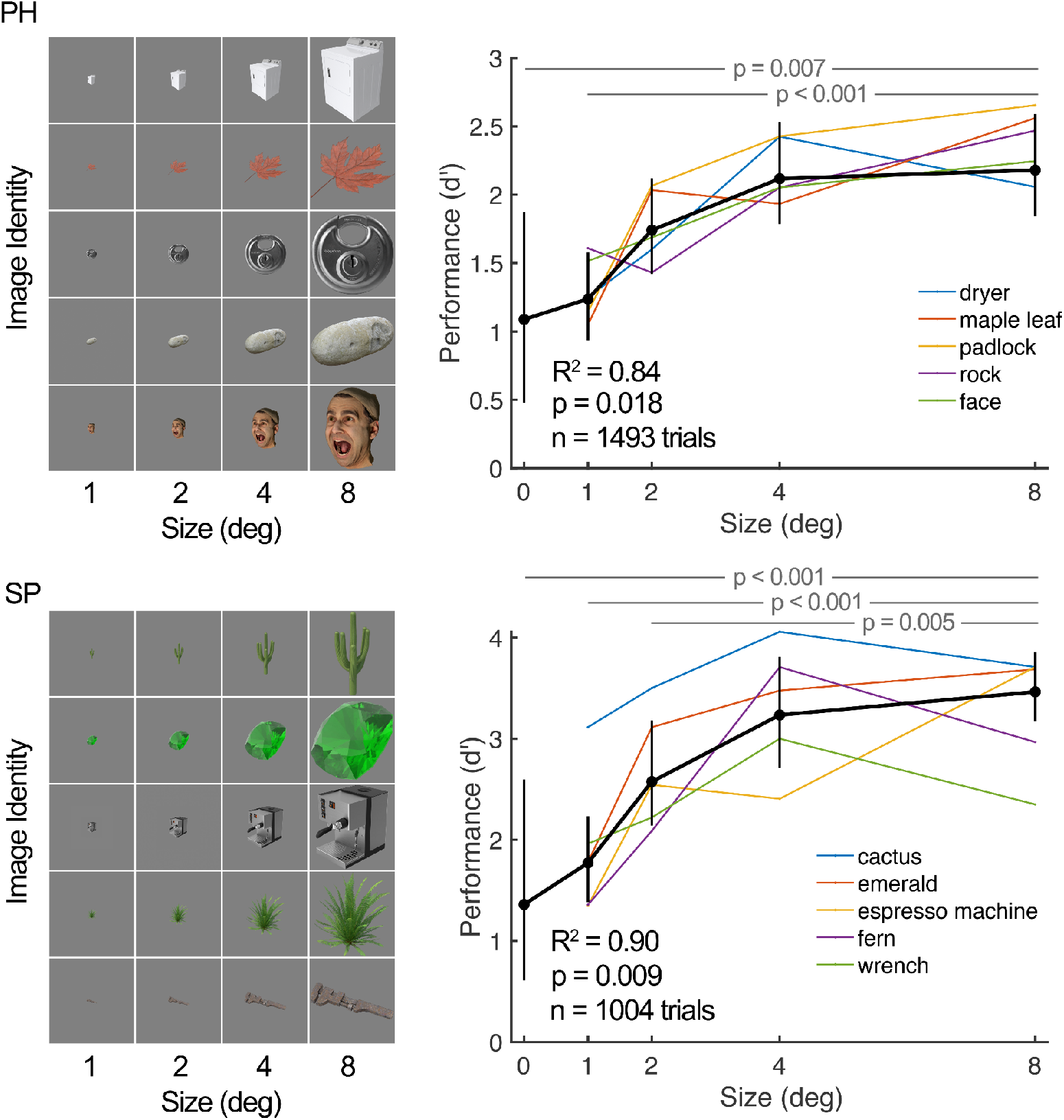
Detectability of optogenetic stimulation of inferior temporal cortex depends on the size of concurrently fixated images. Top: Monkey Ph, Bottom: Monkey Sp. Left: each monkey’s imageset, containing 5 object images and 4 size conditions. The “no image” (uniform gray) condition is not shown. Right: each monkey’s stimulation-detection performance (d’) as a function of image size. The “no image” condition is represented as size 0. Each thin colored line represents data from one object image and the thick black line represents performance across all images. The error bars represent bootstrapped 95% confidence intervals. Image size had a main effect on performance (one-way ANOVA Ph: *F*(3,16) = 18.52, *p* < 0.001; Sp: *F*(3,16) = 5.24, *p* = 0.01). There is a significant correlation between image size and performance (Pearson’s Ph: *r*(20) = 0.82, *p* < 0.001; Sp: *r*(20) = 0.60, *p* = 0.004). Linear regression was performed and a square root curve was used to fit the data (Ph: *R^2^* = 0.84, *t*(3) = 4.73, *p* = 0.018; Sp: *R^2^* = 0.90, *t*(3) = 6.06, *p* = 0.009). The p-values for pairwise comparisons are from post-hoc permutation tests of ANOVA (Benjamini-Hochberg corrected). Note, the face image in Ph’s imageset is computer generated.

## Results

In order to systematically test whether stimulation detectability depends on the size of the concurrently fixated images, the animals performed the optogenetic stimulation-detection task while fixating on randomly presented images of five objects at four size levels (subtending 1, 2, 4, and 8 degrees of visual angle; Figure 2). A ‘no image’ condition was also included. The monkeys’ task was unrelated to the size or content of the images, the animals simply had to report whether a trial did or did not contain cortical stimulation.

Image size had a strong effect on stimulation detectability (Figure 2; one-way ANOVA Ph: *F*(3,16) = 18.52, *p* < 0.001; Sp: *F*(3,16) = 5.24, *p* = 0.01; Pearson’s Ph: *r*(20) = 0.82, *p* < 0.001; Sp: *r*(20) = 0.60, *p* = 0.004). The monkeys’ performance increased with the size of the images; the largest size condition (8 degrees) produced significantly higher performance than the smallest size condition (1 degree) and the no image condition in both animals, as well as the second smallest size condition (2 degrees) in one animal (pairwise comparison permutation tests, followed by Benjamini-Hochberg correction, Ph: *p* < 0.001 for 8 deg vs 1 deg, *p* = 0.007 for 8 deg vs no image; Sp: *p* = 0.005 for 8 deg vs 2 deg, *p* < 0.001 for 8 deg vs 1 deg and 8 deg vs no image).

In a linear regression analysis we used the formula:

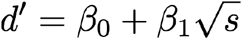

Where *d’* is performance, *s* is image size, *β_0_* is intercept and *β_1_* is slope coefficient. The results confirmed a significant linear relationship between the square root of image size and performance (Ph: *R^2^* = 0.84, *t*(3) = 4.73, *p* = 0.018; Sp: *R^2^* = 0.90, *t*(3) = 6.06, *p* = 0.009). Consistent with our prior results (Azadi, Bohn, Lopez, et al. 2022), this result suggests that the perceptual event evoked by artificial stimulation in IT cortex highly depends on the presence and prominence of the concurrent visual input. Performance on the “no image” condition was still significantly above chance for one animal (permutation test, Ph: *p* = 0.009, Sp: *p* = 0.06), which could be due to the fact that our manipulation did not entirely eliminate the visual input (e.g the monitor was still visible), or a potential residual effect that persists even in the absence of the visual stimulus.

While the animals were required to maintain fixation within a central 2 degree window throughout each trial, regardless of image size, they may have felt compelled to more tightly constrict their fixations on small image trials. If such an asymmetry in fixation effort occured, it could account for the reduced capacity to detect stimulation events for smaller images. To assess whether differences in fixation behavior might account for the observed effect of image size on detection performance, fixational spread was analyzed as a function of image size (Figure 3, left), trial outcome (Figure 3, middle), and detection performance (Figure 3, right). For each trial, the root mean square of all pairwise distances between fixations throughout the trial (measured 1/ms over 1000ms, from image onset until imageset offset) was computed *(fixational spread*). Image size did not have an effect on fixational spread (ANOVA Ph: *F*(3,16), = 0.93 *p* =0.45; Sp: *F*(3,16) = 1.74, *p* = 0.20; Spearman’s Ph: *r*(20) = 0.26, *p* = 0.26; Sp: *r*(20) = 0.09, *p* = 0.70). Fixational spread was also independent of trial outcome (hit, miss, false alarm, correct rejection) (ANOVA Ph: *F*(3,79) = 1.26, *p* = 0.29; Sp: *F*(3,70) = 1.08, *p* = 0.36) and overall performance (Spearman’s Ph: *r*(20) = 0.19, *p* = 0.40; Sp: *r*(20) = 0.05, *p* = 0.83; Ph: *R^2^* = 0.01, *t*(19) = 0.49, *p* = 0.63; *R^2^* = 0.02, *t*(19) = 0.66, *p* = 0.52). Further, the animals did not break fixation more or less often as a function of image size (ANOVA Ph: *F*(3,16) = 0.51, *p* = 0.7; Sp: *F*(3,16) = 0.79, *p* = 0.5; Spearman’s Ph: *r*(20) = 0.05, *p* = 0.8; Sp: *r*(20) = −0.09, *p* = 0.7), or for stimulation trials compared to non-stimulation trials (Ph: *X^2^* (1, *N* = 1775) = 0.003, *p* = 0.96; Sp: *X^2^* (1, *N* = 1438) = 0.11, *p* = 0.74).

**Figure 3.**
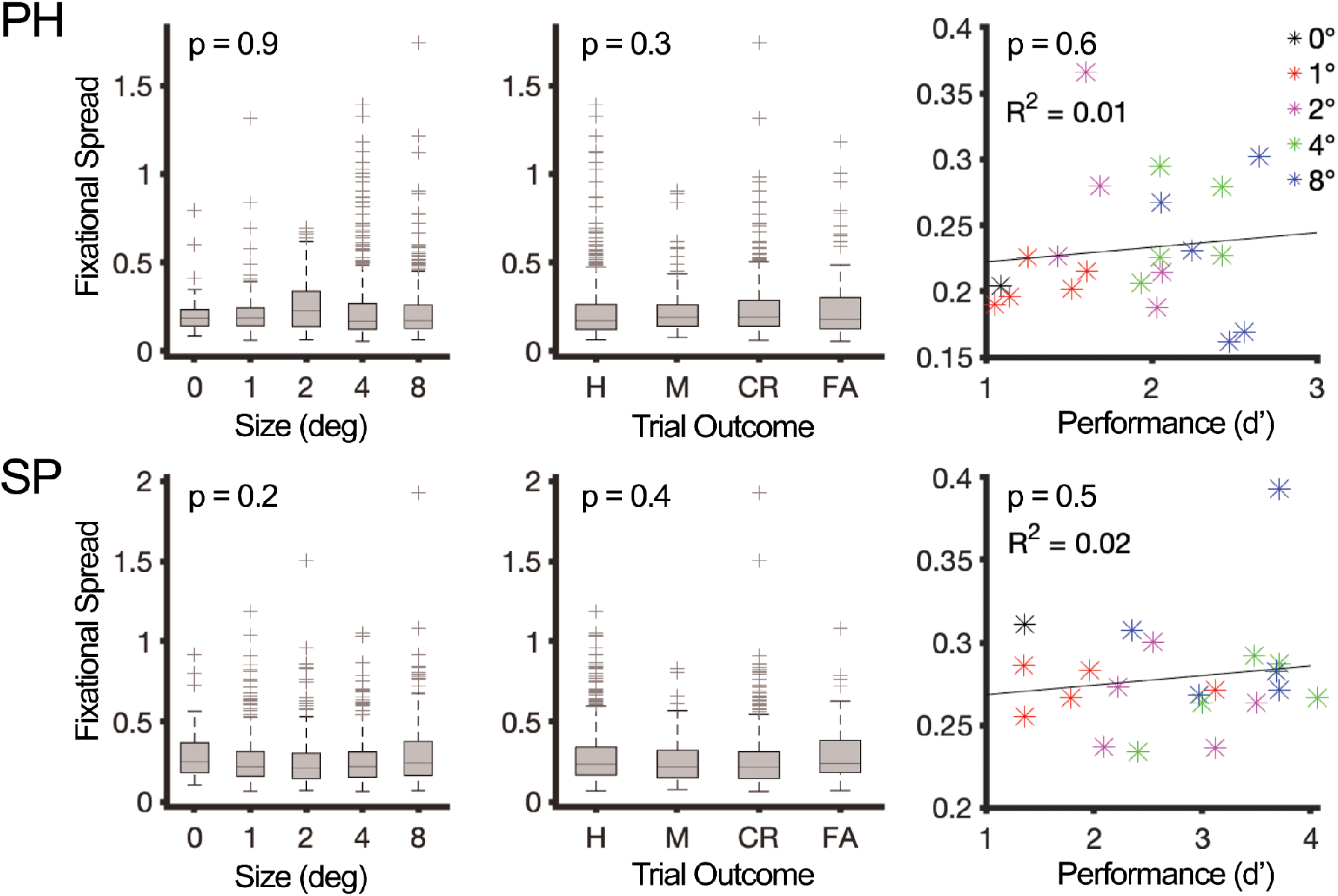
Fixation behavior is independent of image size, trial outcome, and task performance. Top: Monkey Ph, Bottom: Monkey Sp. Left: Fixational spread (RMS of all pairwise distances between fixations throughout each trial, from image onset to image offset) as a function of image size. Crosses represent outliers. Image size did not have an effect on fixational spread (ANOVA Ph: *F*(3,16), = 0.93 *p* =0.45; Sp: *F*(3,16) = 1.74, *p* = 0.20; Spearman’s Ph: *r*(20) = 0.26, *p* = 0.26; Sp: *r*(20) = 0.09, *p* = 0.70). Middle: Fixational spread as a function of trial outcome (Hit, Miss, Correct Rejection, False Alarm). Crosses represent outliers. Fixational spread was independent of trial outcome (ANOVA Ph: *F*(3,79) = 1.26, *p* = 0.29; Sp: *F*(3,70) = 1.08, *p* = 0.36) Right: Fixational spread as a function of stimulation-detection performance (d’). Colored data points represent mean fixational spread for each image (black = no image condition; red = 1 deg; magenta = 2 deg; green = 4 deg; blue = 8 deg). Fixational spread was independent of overall performance (Spearman’s Ph: *r*(20) = 0.19, *p* = 0.40; Sp: *r*(20) = 0.05, *p* = 0.83; Ph: *R^2^* = 0.01, *t*(19) = 0.49, *p* = 0.63; *R^2^* = 0.02, *t*(19) = 0.66, *p* = 0.52).

## Discussion

The present report builds off a foundational study demonstrating that monkeys’ ability to detect identical optogenetic impulses delivered to the same neural population in IT highly depends on what they are looking at during cortical stimulation (Azadi, Bohn, Lopez, et al. 2022). That is, looking at particular images reliably enhanced or impaired stimulation detectability. This observation raised an intriguing question about the phenomenological nature of the perceptual event induced by stimulation: Does stimulation of the same neural population induce a consistent perceptual event (e.g. phosphene), independent of the concurrently fixated image, that is more or less difficult to detect due to figure-ground effects (e.g crowding, contrast)? Or does stimulation induce a variable perceptual event depending on the concurrently fixated image (e.g. distortion)? The prior report addressed this question by diminishing the visibility of the concurrent visual input by reducing the contrast, saturation, and spatial frequency of the concurrently fixated images and found this reduced the monkeys’ ability to detect the stimulation event. The result suggests that the perceptual *nature of the event* evoked by stimulation depends on the concurrent visual input. Put another way, identical stimulation impulses do not elicit identical perceptual events if the concurrent visual input is varied. Consistent with this finding, we show here that image size also has a significant effect on stimulation detectability. Specifically, reducing image size decreases stimulation detectability. That is, smaller images lead to stimulation-induced perceptual events of smaller magnitude.

Why does the concurrent visual input affect stimulation detectability? The definitive answer to this question will require concurrent neural recordings, a key advantage promised by optogenetics over electrical stimulation, nevertheless, the current version of the Opto-Array technology doesn’t yet allow neural recordings. In the meantime, psychophysical studies alone can provide some clues. One possibility is that the visual input modulates the neural thresholds of the stimulated neurons, making the neurons more or less excitable (i.e. perturbable) depending on what the monkey is looking at. The results of the prior visibility experiment and the size experiment presented here are consistent with this idea. While we do not have recordings from the neurons stimulated in this study, prior work has shown that many neurons in IT decrease their firing rate when images are presented at lower contrast (Zoccolan et al. 2007; Rolls and Baylis 1986). As for the case of size, it is traditionally believed that neurons in central IT have very large receptive fields, centered at the fovea, and respond invariant of stimulus size. However, this belief is not well supported by the data and the median size of central IT receptive fields varies dramatically from 26 degrees (Desimone and Gross 1979) to as small as 2.6 degrees degrees (DiCarlo and Maunsell 2003). Moreover, while central IT receptive fields almost always include the fovea, many have peak sensitivity in the parafovea (Op De Beeck and Vogels 2000). In fact, when measured with fMRI, the lateral convexity of IT (where our arrays are implanted) shows a parafoveal bias)(Lafer-Sousa and Conway 2013). Taken together, it is reasonable to assume that larger stimuli have a better chance of engaging IT receptive fields, thus eliciting a larger neural response. Certainly, when there is a stimulus on the screen (as opposed to no image) the targeted neurons are driven to some degree, lowering their effective threshold and making them easier to stimulate. Alternatively, it is possible that artificial stimulation always activates the neurons strongly, thus independently from the visual input, but the downstream read-out mechanisms filter the effects of stimulation differently depending on the state of the rest of the visual system. For example, when there is an object on the screen, the artificially induced activity of a group of IT neurons can be attributed to the onscreen object by the rest of the brain. However, when most of the visual system provides evidence against the existence of an onscreen object (the no image condition), the artificial activation of IT neurons might be ignored by the rest of the brain. This alternative suggests the existence of heuristic mechanisms that gate visual perception. As important as this possibility is, neural recording data is needed to decisively address the question. This may even require more sophisticated models that incorporate readout, feedback, and cortico-cortical dynamics (Jazayeri and Afraz 2017).

At this time there is no off-the-shelf chronically implantable platform capable of simultaneous recording and surface optogenetic stimulation for large nonhuman brains. Technologists should endeavor to develop such tools. fMRI-guided optogenetics could prove a useful interim step. Carrying out functional imaging prior to transducing the cortical tissue and implanting arrays would enable us to direct our optogenetic perturbations to neural populations with specific selectivity profiles, like face patches (Tsao et al. 2003) or color-biased regions (Lafer-Sousa and Conway 2013), and ask whether the types of stimuli those regions prefer enhance detection in the CPD task more so than non-preferred stimuli. We expect the types of stimuli that enhance detection to have common features that relate to the underlying selectivity of the stimulated neurons, as has been observed anecdotally in human patients (Parvizi et al. 2013). An eagle-eyed reader may have noticed that the image sets used in the present work, which were chosen because they were each animal’s top performing objects from an earlier experiment (Azadi, Bohn, Lopez, et al. 2022), had some common features—Sp’s imageset consisted entirely of either green or achromatic objects and Ph’s imageset consisted only of reddish or achromatic objects. Could the perceptual events evoked by stimulation at these sites be chromatic in nature? Answering this and related questions will require additional experiments using many more images and more systematic feature variation. The combined use of the highly sensitive CPD task and the chronically implanted Opto-Array make this undertaking feasible.

The observations presented here have profound implications for the design and interpretation of artificial perturbation studies—one should not assume to get the same psychophysical effect across visual inputs, and perhaps not even the same neural effect. Moreover, these findings raise exciting possibilities and critical considerations for the development of visual prosthetic devices. The traditional approach to visual prosthetics has focused on restoring vision in the profoundly visually impaired by artificial stimulation of the primary visual cortex, where stimulation evokes retinotopic phosphenes. This approach has not been particularly successful and will likely be insufficient to bring about a rich visual experience. How plausible is it to paint a smiling face with phosphenes? Restoring a rich sense of vision will likely require the use of prosthetics applied to IT, where high-level object representations are found. The present work shows first, that IT stimulation induces visual perceptual effects, encouraging its use for prosthetics, and second, that stimulation evoked events in IT depend on the bottom up input. This consequently suggests that if stimulation of IT is considered for visual prosthetics, it should be done in concert with artificial stimulation of low level visual areas. While more experimental and theoretical work is needed in order to explain how local stimulation interacts with the general ongoing activity of the brain, these results expand the horizon of visual prosthetics beyond the primary visual cortex.

## Author contributions

R.L. and K.W. designed the research with guidance from A.A.; R.L. and K.W. performed the research; R.L. analyzed the data; R.A., S.B., E.L., and A.A. performed the surgeries. E.L. and S.B. trained the animals; R.L. prepared the figures; R.L., K.W., and A.A. wrote the manuscript; all authors reviewed the manuscript.

## Data and material availability

The data and material that support the findings of this study are available on request from the corresponding author R.L.

## Acknowledgments

We are grateful to Mark Eldridge for his development of the custom virus-injector array and assistance with the surgeries, to Elia Shahbazi and Timothy Ma for assistance with anatomical reconstructions for surgical planning and useful discussions, and to our funding sources: NIMH Grant ZIAMH002958 and NIMH Intramural Research Training Award (IRTA) Fellowship Program.

